# Subcellular niche segregation of co-obligate symbionts in whiteflies

**DOI:** 10.1101/2022.11.18.517168

**Authors:** Akiko Fujiwara, Xian-Ying Meng, Yoichi Kamagata, Tsutomu Tsuchida

## Abstract

Many insects contain endosymbiotic bacteria within their bodies. In multiple endosymbiotic systems comprising two or more symbionts, each of the symbionts is generally localized in a different host cell or tissue. *Bemisia tabaci* (Sweet potato whitefly)possesses a unique endosymbiotic system where co-obligate symbionts are localized in the same bacteriocytes. Using fluorescence *in situ* hybridization, we found that endosymbionts in *B. tabaci* MEAM1 occupy distinct subcellular habitats, or niches, within a single bacteriocyte. *Hamiltonella* was located adjacent to the nucleus of the bacteriocyte, while *Portiera* was present in the cytoplasm surrounding *Hamiltonella*. Immunohistochemical analysis revealed that the endoplasmic reticulum separates the two symbionts. Habitat segregation was maintained for longer durations in female bacteriocytes. The same segregation was observed in three genetically distinct *B. tabaci* groups (MEAM1, MED Q1, and Asia II 6) and *Trialeurodes vaporariorum*, which shared a common ancestor with *Bemisia* over 80 million years ago, even though the coexisting symbionts and the size of bacteriocytes were different. These results suggest that the habitat segregation system existed in the common ancestor and was conserved in both lineages, despite different bacterial partners coexisting with *Portiera*. Our findings provide insights into the evolution and maintenance of complex endosymbiotic systems and highlight the importance of organelles for the construction of separate niches for endosymbionts.

**Importance:** Co-obligate endosymbionts in *B. tabaci* are exceptionally localized within the same bacteriocyte (a specialized cell for endosymbiosis), but the underlying mechanism for their coexistence remains largely unknown. This study provides evidence for niche segregation at the subcellular level between the two symbionts. We showed that the endoplasmic reticulum is a physical barrier separating the two species. Despite differences in co-obligate partners, this subcellular niche segregation was conserved across various whitefly species. The physical proximity of symbionts may enable the efficient biosynthesis of essential nutrients via shared metabolic pathways. The expression “Good fences make good neighbors” appears to be true for insect endosymbiotic systems.

## Introduction

Numerous insects contain endosymbiotic bacteria within their bodies. Some endosymbionts are obligate, with crucial roles in host growth and reproduction, by providing essential nutrients, while others are facultative (1, 2). In multiple endosymbiotic systems comprising two or more symbionts, each of the symbionts is generally localized in a different host cell or tissue. This type of localization is observed in various insect species, such as aphids, spittlebugs, leafhoppers, scale insects, adelgids, psyllids, and cicadas (1, 3–13) (*Supporting data only for review* in Supplemental Information). The compartmentalization of multiple symbionts into different host cells is considered as an important step in evolution, to reduce direct conflict between the multiple symbionts and control them within the same host (12, 14).

The sweet potato whitefly *Bemisia tabaci* (Hemiptera: Aleyrodidae) is a cryptic species complex comprising more than 44 genetic groups based on the mitochondria cytochrome oxidase I sequences (15). Among the genetic groups, Middle East Asia Minor 1 (MEAM1) and Mediterranean Q1 (MED Q1) are globally important pests, and both possess two phylogenetically distinct types of endosymbiotic bacterium, *Candidatus* Portiera aleyrodidarum and *Candidatus* Hamiltonella defense (hereinafter called *Portiera* and *Hamiltonella*, respectively). They are co-obligate symbionts based on the virtually universal infection observed in natural populations (16–23), the decrease in host fitness caused by symbiont elimination (24, 25), and their genome contents for producing essential amino acids and cofactors (26–29), although mathematical metabolic models assuming limited conditions suggested that *Hamiltonella* shows a nutritionally parasitic state (30).

Interestingly, previous studies have reported that these symbionts in *B. tabaci* are co-localized in the same bacteriocytes (31–33), suggesting possible competition between the symbionts for limited space and resources. Nevertheless, *Portiera* and *Hamiltonella* populations are large and exhibit synchronous dynamics in MEAM1 (34). These data suggest that some mechanism exist to maintain the two essential symbionts in the same bacteriocytes. One mechanism for avoiding conflict between the symbionts would be niche segregation, although it has never been reported at the subcellular level. In this study, we demonstrated that habitats of the symbionts are segregated within a single bacteriocyte by the endoplasmic reticulum (ER). Our results also suggest that this segregation system operates not only in specific genetic groups of *B. tabaci* but also in other whitefly species.

## Results and Discussion

### Temporal dynamics of co-obligate symbionts

We used quantitative polymerase chain reaction (PCR) to investigate the population dynamics of the two symbiotic bacteria *Portiera* and *Hamiltonella* in MEAM1. Both populations increased during nymphal growth, peaked in actively reproducing young adults, and declined in older whiteflies (Fig. S1). The primary role of these symbionts is to provide nutrients to the host (26–29). Consistent with that role, their population dynamics correspond to the necessity for the host growth and reproduction, as reported for other obligate symbionts of multiple insect hosts, including the pea aphid (35–37), cereal weevils (38), and a different MEAM1 strain (34). The populations of both symbionts in females were maintained at relatively higher levels for longer durations than those for the male populations (Fig. S1). These results suggest that the symbiont populations are differentially regulated depending on the host’s sex.

### *Portiera* and *Hamiltonella* occupy the same bacteriocytes but segregate into different subcellular niches

Previous studies using FISH analysis (39, 40) suggested that *Hamiltonella* is distributed around the nucleus of the bacteriocyte while *Portiera* is distributed in the cytoplasmic regions. However, the distribution pattern was not clear as the observations were made on a single slice of the bacteriocyte. Therefore, we acquired 3D spatial data depicting the distribution of the two symbionts in bacteriocytes using Z stack analysis. Confocal laser scanning microscopy of the teneral adults of MEAM1 revealed distinct niche for the two symbionts within the same bacteriocytes: *Hamiltonella* was located adjacent to the nucleus of the bacteriocyte, while *Portiera* was located in the cytoplasm, surrounding *Hamiltonella* (Fig. 1A, Movie S1). In contrast, *Rickettsia*, which generally infects MEAM1 as a facultative symbiont (16, 19–23, 41, 42), were scattered all over the body and rarely observed in the bacteriocytes (Fig. S2*A* and *B*).

**Figure 1.**
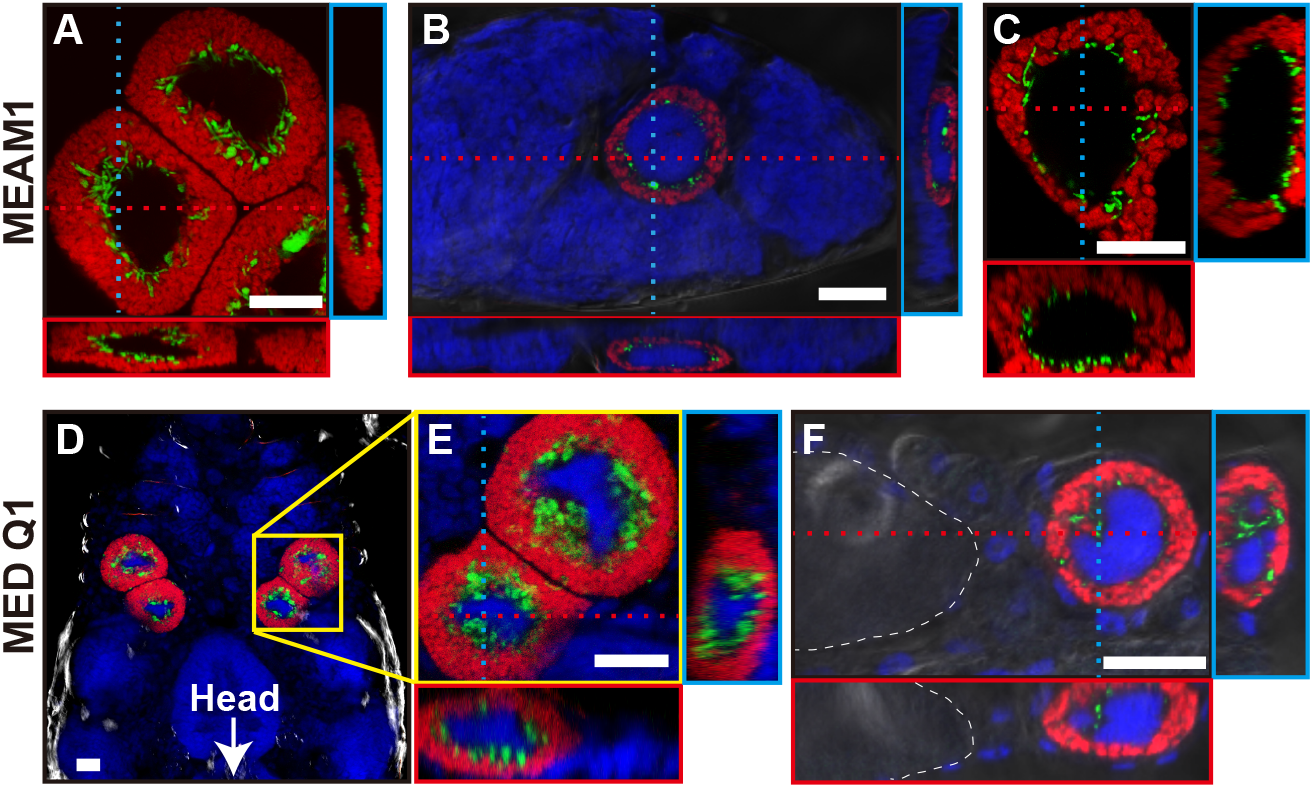
Localization of *Portiera* (red) and *Hamiltonella* (green) in bacteriocytes of *Bemisia tabaci*. (*A–C*) MEAM1: (*A*) bacteriocytes dissected from an adult female, (*B*) A 3-day-old egg, and (*C*) a bacteriocyte dissected from a fourth-instar nymph. (*D–F*) MED Q1 strain: (*D*) Bacteriocytes of the fourth-instar nymph, (*E*) enlarged image of the regions indicated by a yellow square in (*D*), and (*F*) a bacteriocyte just before entering the egg in a teneral adult female. In (*A–C, E*, and *F*), orthogonal views of Z-stack images are shown; red and blue dashed lines indicate corresponding points in the orthogonal planes. In (*B, D, E*, and *F*), host nuclear DNA is visualized in blue. In (*F*), white dashed lines indicate the outline of the egg. Bars, 20 µm.

In *B. tabaci*, an intact bacteriocyte-bearing symbiont is transferred to each egg in adult females (43) that persists through embryogenesis (44). Niche segregation between *Portiera* and *Hamiltonella* was observed in the bacteriocytes transferred into the eggs (Fig. 1*B*) of MEAM1 and was maintained from nymphs (Fig. 1*C*) to female and male young adults (Fig. S3*A* and *B*), during which the metabolic function of the symbionts becomes necessary for host growth and reproduction. At 15 d after adult eclosion, the observed habitat segregation was disrupted only in male bacteriocytes (Fig. S3*C* and *D*). This observation was consistent with the report that apoptosis and autophagy occurred in male bacteriocytes but not in females (33). *Portiera* and *Hamiltonella* titers remarkably declined only in males at this stage (Fig. S1); hence it is conceivable that the long-term stable habitat in female bacteriocytes evolved to ensure the vertical transmission of both symbionts.

### The endoplasmic reticulum segregates symbionts within bacteriocytes

We performed electron microscopy to further understand the mechanisms underlying the subcellular niche segregation of symbionts in bacteriocytes. The results revealed the presence of rod-shaped bacteria next to the nucleus and unstructured hypertrophic bacteria on the outer cytoplasm (Fig. 2*A*). The shapes and locations of the two bacteria types were consistent with those of *Hamiltonella* and *Portiera*, as demonstrated by FISH (Fig. 1*A*–*C*). Interestingly, an ER-like multiple membrane structure was observed around *Hamiltonella* (Fig. 2*B*).

**Figure 2.**
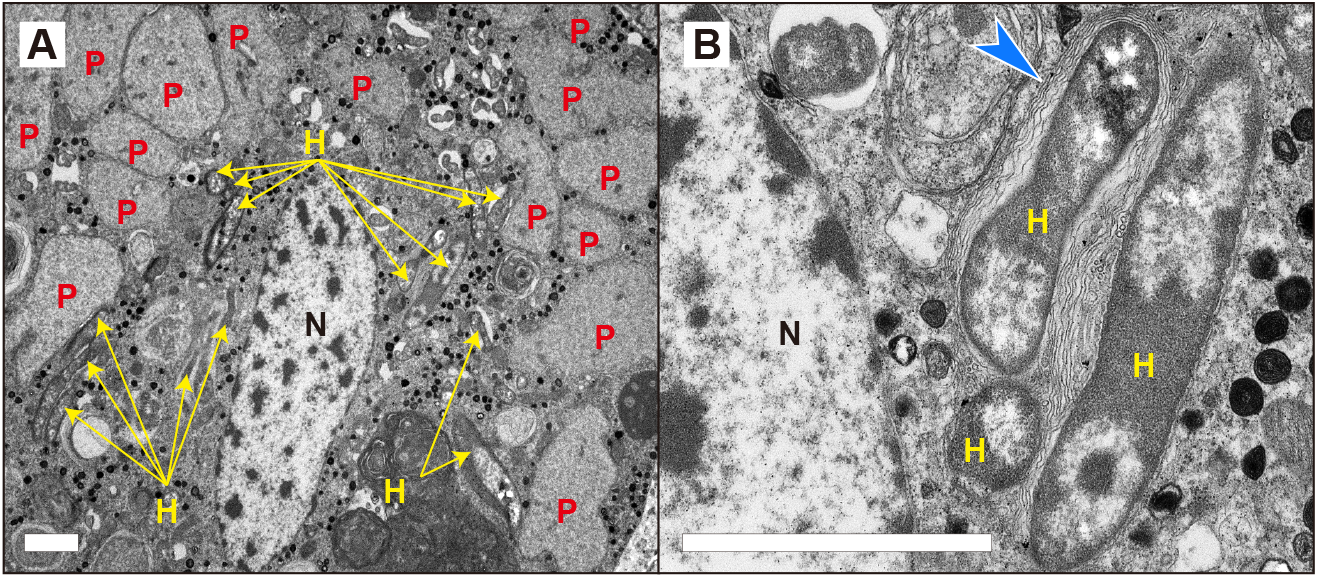
Transmission electron micrographs (TEM) of a bacteriocyte in *B. tabaci* MEAM1. (*A*) Unstructured hypertrophic bacteria (*Portiera*) in the cytoplasm and rod-shaped bacteria (*Hamiltonella*) around the nucleus of the bacteriocyte. (*B*) Enlarged image of *Hamiltonella*. N, nucleus of a bacteriocyte; P, *Portiera*; H, *Hamiltonella*. Blue arrowhead, ER-like structures surrounding *Hamiltonella*. Bars, 2 µm.

The ER and *Hamiltonella* were simultaneously detected using a combination of immunohistochemistry and FISH at the young adult stage. Strong fluorescence signals for the ER marker were detected around the nuclei of bacteriocytes and surrounded the fluorescence signals for *Hamiltonella* (Fig. 3*A* and Fig. S4*A*). A three-dimensional analysis using composite Z-stack images revealed that the ER encompassed *Hamiltonella* (Fig. 3*A*). These results confirmed that the ER partitioned *Hamiltonella* and *Portiera*. At 15 d after adult eclosion, the ER and cell structure disruption was observed in males (Fig. S4*C*), possibly due to programmed cell death (33). In comparison, the ER was detected between the co-obligate symbionts in females (Fig. S4*B*). These results indicate that the ER-mediated subcellular niche segregation of the symbionts lasts longer in females. It suggests that the ER has an important role in maintenance of stable niche for each symbiont especially in female bacteriocytes which are needed to be transferred to the next generation (44).

**Figure 3.**
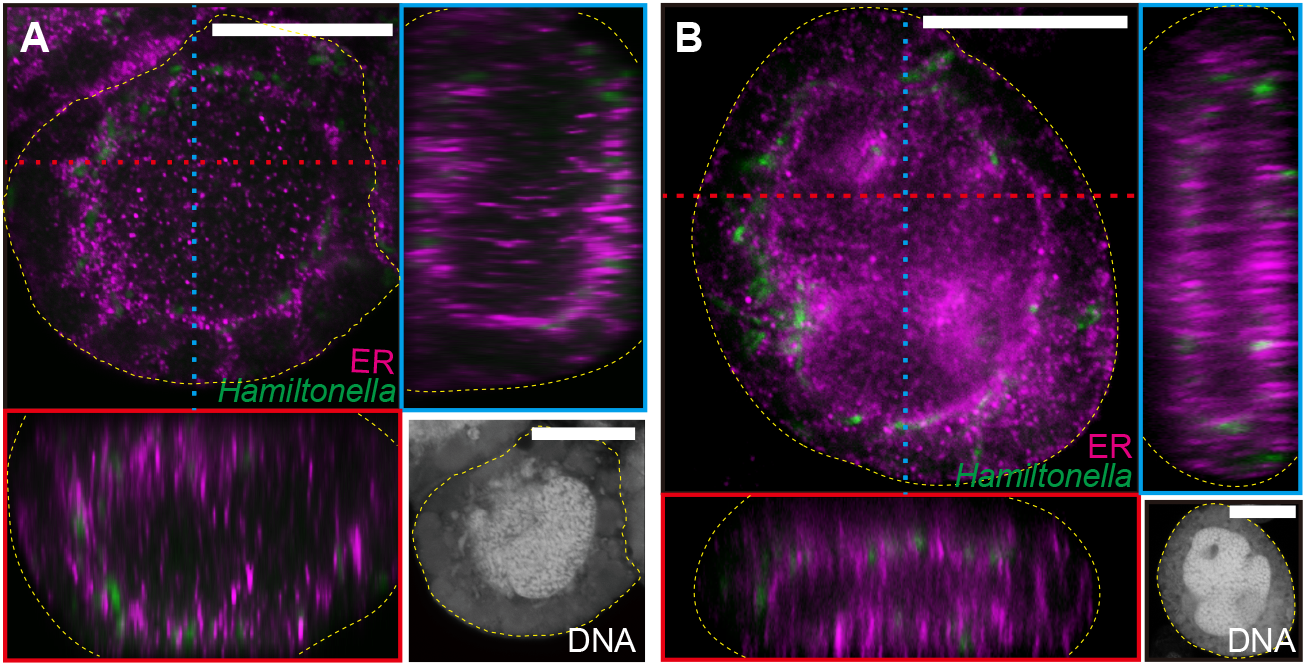
Localization of *Hamiltonella* and the ER in a bacteriocyte of a young adult female (1–5 d after eclosion) of MEAM1(*A*) and MED Q1(*B*). Orthogonal views of Z-stack images are shown. Red and blue dashed lines indicate corresponding points in the orthogonal planes. *Hamiltonella* and ER are shown in green and violet, respectively. Panels in the bottom right corner of each figure are DAPI-stained images, showing nuclei in the center of the bacteriocytes and *Portiera* and *Hamiltonella* around the nuclei. Yellow dashed lines indicate the outline of the bacteriocytes. Bars, 20 µm.

### Subcellular niche segregation in other *B. tabaci* subgroups and more distantly related species

The *B. tabaci* subgroup MED Q1 also possesses *Portiera* and *Hamiltonella* as co-obligate symbionts (16-21, 26). FISH analysis indicated that *Portiera* and *Hamiltonella* are segregated within individual bacteriocytes through earlier developmental stages in MED Q1 (Fig. 1*D*–*F*), as observed in MEAM1 (Fig. 1*A*–*C*). The facultative symbiont *Cardinium* in MED Q1 did not show specific localization in bacteriocytes (Fig. S2*C* and *D*). Combining FISH and immunohistochemistry, the ER membrane encompassed *Hamiltonella* and separated it from *Portiera* (Fig. 3*B* and Fig. S4*D*). Cytological staining of the dissected living bacteriocytes from young adults also demonstrated that *Hamiltonella* was sterically surrounded by the ER and was segregated from *Portiera* (Fig. S5).

Furthermore, we assayed for subcellular habitat segregation in a distantly related genetic group of *B. tabaci*, Asia II 6. Asia II group is frequently infected with *Candidatus* Arsenophonus sp. (hereafter called *Arsenophonus*) in addition to *Portiera* but not with *Hamiltonella* (19, 20, 45). Using FISH, we demonstrated that *Arsenophonus* is located adjacent to the nucleus of the bacteriocyte in both adults and developing eggs (Fig. 4). The localization pattern of *Arsenophonus* in Asia II 6 was identical to that of *Hamiltonella* in both MEAM1 (Fig. 1*A*–*C*) and MED Q1 (Fig. 1*D*–*F*). Moreover, *Arsenophonus* around the nucleus was surrounded by the ER (Fig. S4*E*), similar to that of *Hamiltonella* in MEAM1 and MED Q1 (Fig. S4*A, B* and *D*).

**Figure 4.**
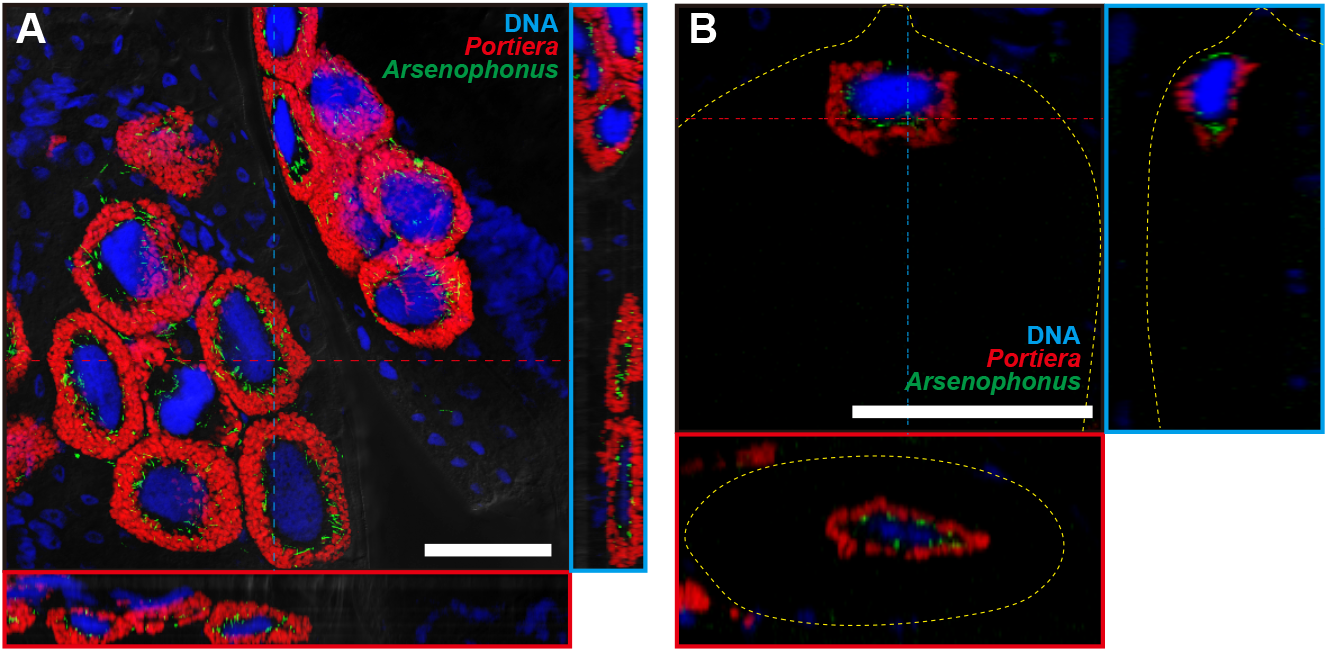
Localization of *Portiera* (red) and *Arsenophonus* (green) in a putative young adult female *B. tabaci* Asia II 6. (*A*) Bacteriocytes in the abdomen. (*B*) developing egg within the female. Orthogonal views of Z-stack images are shown. Red and blue dashed lines indicate corresponding points in the orthogonal planes. Host nuclear DNA is visualized in blue. In (*B*), yellow dashed lines indicate the outline of the egg. Bars, 50 µm.

*Trialeurodes vaporariorum* belongs to the same family as *B. tabaci, Aleyrodinae* (46). *Arsenophonus* has been detected at high frequencies in *T. vaporariorum* (22, 32, 47, 48) and is considered an essential symbiont, supplying or complementing the nutrients that are not produced by coexisting *Portiera* (49). FISH revealed that *Portiera* and *Arsenophonus* also exhibit subcellular habitat segregation in the same bacteriocytes of *T. vaporariorum* (Fig. 5*A*). Immunohistochemical staining with FISH indicated that the ER encloses *Arsenophonus*, separating it from *Portiera* (Fig. 5*B* and Fig. S4*F*). Bacteriocytes in *T. vaporariorum* (ca. 20 µm in diameter) are considerably smaller than those of *B. tabaci* (ca. 30 to 50 µm). Despite the morphological and phylogenetic differences, the same system in bacteriocytes is involved in symbiont niche partitioning. This suggests that the subcellular niche segregation of the symbionts evolved in the common ancestor and have been conserved to the present.

**Figure 5.**
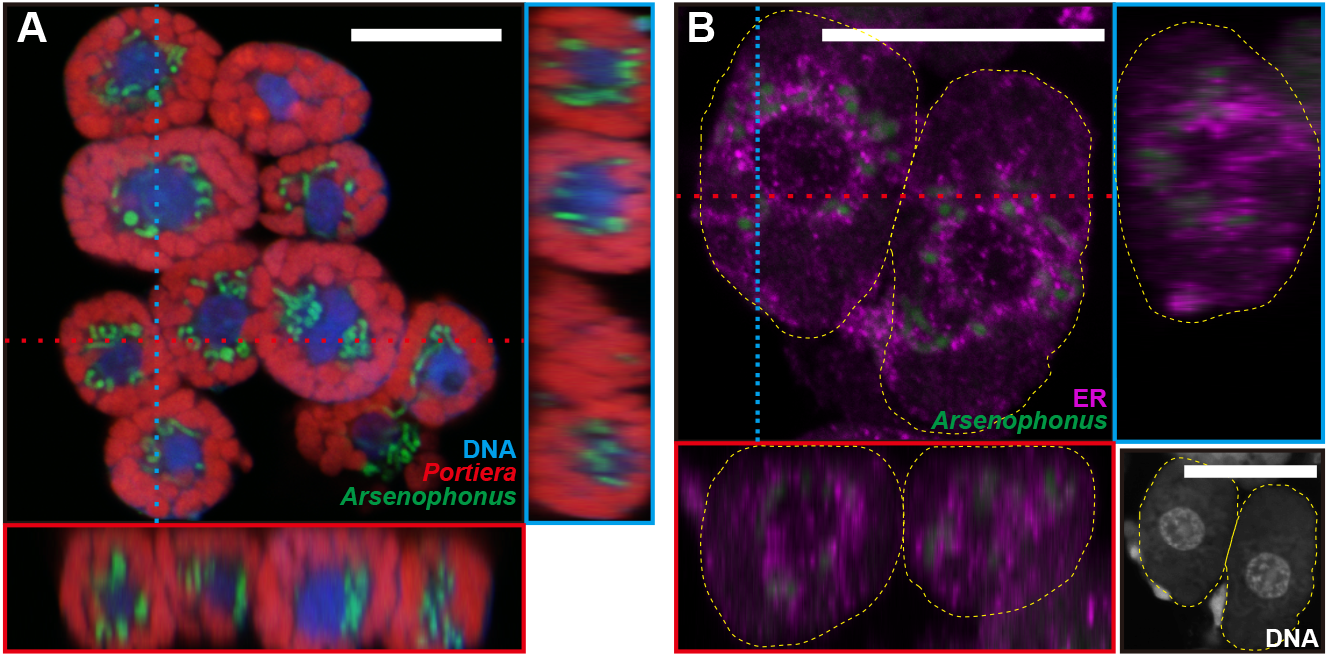
Localization of *Portiera* (red), *Arsenophonus* (green), and the ER (violet) in bacteriocytes of *Trialeurodes vaporariorum*. (*A*) FISH image and (*B*) immunohistochemistry combined with FISH (*B*). Orthogonal views of Z-stack images are shown. Red and blue dashed lines indicate corresponding points in the orthogonal planes. In (*B*), panel in the bottom right corner is DAPI-stained images and yellow dashed lines indicate the outline of a bacteriocyte. Bars, 20 µm.

### Biological implications, evolution, and possible mechanisms underlying subcellular symbiont segregation

Previous ecological studies have shown that niche segregation reduces the strength of competition between species that exploit the same limiting resource (50–52). Niche segregation is found in various taxa, including endosymbiotic bacteria in insects, in which each endosymbiont generally localizes in different host cells or organs (1, 3–12). The compartmentalization of multiple symbionts into different host cells is considered as an important step in the evolution to reduce direct conflict between the multiple symbionts and control them within the same host (14). The endosymbiotic system in *B. tabaci* is a rare exception because co-obligate symbionts occupy the same bacteriocytes (31, 32). In this study, we revealed that the endosymbionts in *B. tabaci* and *T. vaporariorum* are separated by the ER and show niche segregation at the subcellular level. This novel finding indicates that the niche segregation of endosymbionts can be established not only in different host cells but also within a single cell.

It is intriguing why the symbiont co-habitation has evolved exceptionally in the whiteflies. In certain whitefly species, including *B. tabaci* and *T. vaporariorum*, one or more entire bacteriocytes bearing symbionts are transferred to the developing egg (1, 43, 44, 53). Under the unique transmission manner, one of the symbiont partners could be lost by the failure of transmission if the two co-obligate symbiotic bacteria were separated into different cells. Hence, it is conceivable that the unique transmission machinery of the symbionts has driven the evolution of the symbiont co-localization in whiteflies. To clarify the evolutionary process of co-habitation and subcellular niche segregation of the symbionts, further studies are required on the vertical transmission and localization manners of the symbionts in various whitefly species.

In some co-obligate symbiotic systems in insects, each symbiont can complete the biosynthetic pathway of some essential nutrients, such as essential amino acids or vitamins, on its own or with host genes (13, 54–58). In other co-symbiotic systems, metabolic interdependence for some essential nutrients is present, wherein one of the symbionts possesses genes necessary for the first part of the biosynthetic pathway, and the other symbiont has genes necessary for the latter part of the pathway (10, 12, 13, 59–60). In such a system, metabolic intermediates must diffuse or be translocated from one symbiont to another to produce essential nutrients in adjacent host cells. However, the co-obligate symbiotic system in *B. tabaci* has a more complex interdependence; enzymes for essential amino acids (*e*.*g*., lysine) are dispersed across both *Portiera* and *Hamiltonella*, and metabolic intermediates should be transported between the symbionts multiple times for production (26–28) (Fig. S6*A*). Although several metabolic duplications are present in the host genome (27, 28, 61), the physical proximity between *Portiera* and *Hamiltonella* in the same bacteriocyte, which is stably maintained during development (Fig. 1), may be adaptive for the efficient production of essential amino acids using intertwined biosynthetic pathways (Fig. S6*C*). It should be noted that mealybugs also have intertwined metabolic pathways in their co-obligate symbiotic systems; gamma-proteobacteria are located inside the beta-proteobacterium *Tremblaya*, and metabolic intermediates are transported between them multiple times to produce nutrients (62–65). Accordingly, reducing the distance between symbionts could be a driving force for the evolution of complex interdependent biosynthetic pathways.

*B. tabaci* is a cryptic species complex with reportedly more than 44 genetic groups (15). In this study, we found the subcellular habitat segregation of two symbionts in three distinct genetic groups in *B. tabaci*, MEAM1, MED Q1, and Asia II 6 (Fig. 1, 4 and Fig. S4*A*-*E*). Moreover, the same localization pattern of symbionts was detected in *T. vaporariorum* (Fig. 5 and Fig. S4*F*) belonging to the different genera. In Asia II 6 and *T. vaporariorum*, the subcellular niche occupied by *Hamiltonella* in MEAM1 and MED Q1 was occupied by *Arsenophonus*. Similar to *Hamiltonella* in *B. tabaci*, it has been suggested that *Arsenophonus* shares metabolic pathways for essential amino-acid synthesis with *Portiera* and the host *T. vaporariorum* (Fig. S6*B*) (49, 66). *Arsenophonus* might establish co-obligate symbiotic systems with *Portiera* in Asia II 6 as well as in *T*.

*vaporariorum*. In many geographic regions, *Hamiltonella* and *Arsenophonus* are rarely found in the same individual of *B. tabaci* and *T. vaporariorum* (16-23, 67), suggesting that the two symbiotic bacteria are competing over functional and cytological niches. Moreover, it appears that the habitat segregation system was established in a common ancestor of species in *Aleyrodinae* and was conserved, while the bacterial partners coexisting with *Portiera* have been replaced (Fig. 6). To understand the generality and the evolutionary origin of the habitat segregation system, detailed analyses of other genetic groups in *B. tabaci* and diverse whitefly species are required.

**Figure 6.**
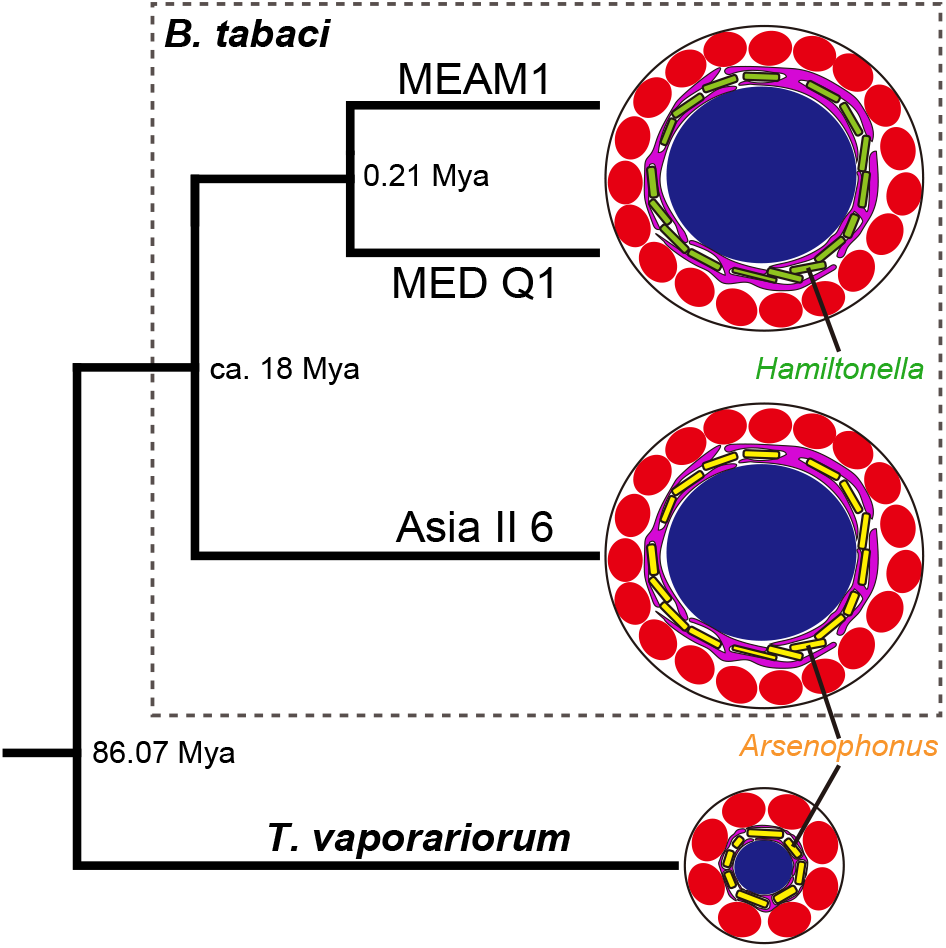
Phylogenetic relationships among whiteflies and their symbiotic system in bacteriocytes. Numbers at internal nodes indicate the divergence date (Mya: million years ago) estimated by Santos-Garcia *et al*. (66). The size of bacteriocytes is shown on the same scale.

As mentioned above, the bacterial partner coexisting with *Portiera* in bacteriocytes varied depending on the species and groups of host insects. This indicates that some molecular and cellular mechanisms underlying subcellular habitat segregation are mainly attributed to the host insects. It is also possible that the symbionts in whiteflies have some properties associated with the ER in bacteriocytes. It is noteworthy that human pathogenic bacteria, such as *Legionella pneumophila, Brucella abortus*, and *Simkania negevensis*, exploit the host ER to generate a niche for replication (68–71). Recent studies have revealed that the organelles have an intimate association with endosymbiotic bacteria and play key roles in the maintenance and control of the endosymbiotic system in insects (72–76). This study provides novel insight into the interaction between symbionts and organelles. To clarify the molecular mechanisms underlying subcellular niche segregation, further studies are required from the perspectives of both the host and the symbionts. Super-resolution microscopy and cryo-electron microscopy will be useful in revealing details about the sites of interaction between the symbionts and ER and inferring molecular mechanisms.

## Materials and methods

### Materials

Whitefly strains and their symbionts used in the study are listed in Table S1. *Bemisia tabaci* was maintained on cabbage (*Brassica oleracea*) and *Trialeurodes vaporariorum* on cucumber (*Cucumis sativus*) at 25 ± 1°C and 40–60% relative humidity in a long-day regimen (16L: 8D). Adult *B. tabaci* Asia II 6 insects were collected from the field in May 2017 and immediately stored in 100% acetone until use (77). Additional detailed information is provided in Table S1 and Fig. S7.

### DNA extraction and quantitative PCR analysis of symbionts

Ten individuals of MEAM1 at each developmental stage (Fig. S7), from egg to adult, were collected. Eggs and nymphs were collected without distinction between males and females as the sex of whiteflies is indistinguishable at immature stages. In the adult stages, whiteflies from both sexes were separately collected. Samples were preserved in 100% acetone (77) until DNA extraction. The total genomic DNA of the insect individual and its symbionts was extracted using the NucleoSpin Tissue XS Kit (MACHEREY-NAGEL, Duren, Germany), according to the manufacturer’s instructions. From the eggs, the DNA was extracted from groups of 10 individuals. *Portiera* and *Hamiltonella* were quantified in terms of 16S ribosomal RNA gene copies using the Mx3005P system (Agilent Technologies, Santa Clara, CA, USA) with THUNDERBIRD SYBR qPCR Mix (Toyobo, Osaka, Japan), and specific primer sets (Table S2). PCR conditions were: 95°C for 3 min, followed by 40 cycles of 95°C for 30 s and 55°C for 30 s, and a final extension at 72°C for 30 s. The quantitative PCR analysis was conducted by using a standard curve method, as described previously (78).

### Fluorescence *in situ* hybridization (FISH)

To examine the localization of the symbionts, the insect whole-body or dissected tissue specimens were fixed in Carnoy’s solution (EtOH: chloroform: glacial acetic acid, 6:3:1), bleached in 6% hydrogen peroxide in EtOH, and subjected to whole-mount FISH, as described (79). Fluorochrome-labelled oligonucleotide probes are listed in Table S2. Host-cell nuclei were counterstained with 4,6-diamino-2-phenylindole (DAPI). The specificity of *in situ* hybridization was confirmed using a no-probe control and an RNase digestion control (78).

### Transmission electron microscopy

Twenty teneral adults of MEAM1 (1 d old after eclosion) were dissected in 2.5% glutaraldehyde in 0.1 M phosphate buffer (pH 7.4), and the dissected bacteriocytes were pre-fixed with the fixative at 4°C overnight. Subsequently, the samples were post-fixed with 2% osmium tetroxide in 0.1 M phosphate buffer (pH 7.4) at 4°C for 90 min before dehydration with a graded ethanol series. The dehydrated specimens were embedded in EPON 812 resin, processed into ultrathin sections (approximately 80 nm thick) using an ultramicrotome EM UC7 (Leica, Wetzlar, Germany), mounted on copper meshes, stained with uranyl acetate and lead citrate, and observed under a transmission electron microscope (80kV; H-7600 Hitachi, Tokyo, Japan).

### Simultaneous detection of symbionts and endoplasmic reticulum

Ten to twenty mixed-sex individuals of 1*–*5 d or 15 d after adult eclosion were collected from the laboratory strains, *B. tabaci* MEAM1 and MED Q1 and *T. vaporariorum* (Table S1). For *B. tabaci* Asia II 6, acetone-preserved samples were used owing to the difficulty of obtaining fresh materials. After samples were washed with 70% EtOH, bacteriocytes were collected by dissection in phosphate-buffered saline (PBS). The bacteriocytes were transferred to 6.5-mm Transwell dishes with 8-µm pore polycarbonate membrane inserts (Corning, New York, NY, USA) and fixed in 4% paraformaldehyde (PFA in phosphate buffer) for 3 h at 25°C. After fixation, bacteriocytes were washed thrice in PBS with 0.3% TritonX-100 (PBSTx) for 30 min and then soaked thrice in hybridization solution (20 mM Tris-HCl pH 8.0, 0.9 M NaCl, 0.01% SDS and 30% formamide) for 5 min. Then, fluorochrome-labeled probes were hybridized overnight at 25°C. The oligonucleotide probes (Table S2) were used for *Portiera* detection. For *Hamiltonella* or *Arsenophonus* detection, Stellaris RNA FISH probe sets were used (Biosearch Technologies, Novato, CA, USA) (Table S2) as previously described (80). Host-cell nuclei were counterstained with DAPI during hybridization. Subsequently, the samples were refixed in 4% PFA for 7 h at 4°C as described (81) with minor modifications. Then, bacteriocytes were washed thrice in PBSTx and blocked with 1% gelatin in PBS for 30 min at 25°C. After blocking, bacteriocytes were incubated with the KDEL ER marker antibody (10C3) (Santa Cruz, Dallas, TX, USA; cat. no. sc-58774) diluted (1:50) in PBS containing 1% gelatin and 0.05% Tween 20 overnight at 4°C. Bacteriocytes were washed thrice in PBSTx and then incubated with Alexa Fluor Plus 555-or 647-conjugated goat anti-mouse IgG secondary antibody (Thermo Fisher Scientific, Waltham, MA, USA) at a dilution of 1:100 for 3 h at 25°C.

### Laser confocal microscopy

Samples were mounted with ProLong Diamond (Thermo Fisher Scientific). Images were obtained using a LSM5 Pascal or LSM710 microscope and analyzed using LSM5 Pascal Image and LSM ZEN2009 software (Carl Zeiss, Oberkochen, Germany).

### Cytological staining

Staining bacteriocytes was performed as described previously (82), with some modifications. Bacteriocytes were dissected from female *B. tabaci* MED Q1 (Table S1) within 3 d after adult eclosion in Buffer A (35 mM Tris–HCl, pH 7.6, containing 10 mM MgCl_2_, 25 mM KCl, and 250 mM sucrose) and stained with 4 µM SYTO16 (Thermo Fisher Scientific, Waltham, MA, USA) for nucleic acids (bacteria and host) and 4 µM ER-Tracker Red (Thermo Fisher Scientific) for ER in Buffer A for 2 h at 37°C on the SkyLight Glass Base Dish 3971-035-SK (IWAKI, Shizuoka, Japan). Images were obtained using a laser confocal microscope (LSM710) and analyzed using the LSM software ZEN2009 (Carl Zeiss, Oberkochen, Germany).

### Statistics

To evaluate differences in the symbiont titers between females and males at each adult stage (days 1, 15 and 30), the Mann-Whitney U test with Bonferroni’s correction was adopted. Analyses were conducted with R v.3.5.3 software (http://www.r-project.org).

## Acknowledgements

We thank Y. Horita, D. Hwang, K. Moronaga, N. Murakami, Y. Utsuno, and M. Watanabe for technical assistance; N. Haruyama, K. Kijima, T. Kitamura, J. Ohnishi, I. Ohta, and T. Uesato for providing insect samples; and S. Egoshi, K. Dodo and M. Yoshida for useful comments. Part of this study was supported by JSPS KAKENHI Grant Number 18K05673 (to T.T.) and grants from the Project of the NARO Bio-oriented Technology Research Advancement Institution (Research program on the development of innovative technology). A.F. was supported by grants from the RIKEN Special Postdoctoral Researcher (SPDR) Program and Leading Initiative for Excellent Young Researchers (LEADER) Program.

## Author contributions

A.F. and T.T. designed research; A.F. and Y.M. performed research; Y.K. contributed new analytic tools; A.F. and T.T. analyzed data and wrote the paper.

## Supplemental Movie Legend

Movie S1. Z-stack images of bacteriocytes harboring *Portiera* (red) and *Hamiltonella* (green) in *B. tabaci* MEAM1. Bar, 50 µm. In total, 55 spatially consecutive images were collected by confocal microscopy (depth interval = 0.5 µm) and were processed using Zeiss LSM5 Pascal Image software. Nuclei in the center of bacteriocytes are not shown in the movie to clarify the localization patterns of the two symbionts.

## Supplemental Figure Legends

Fig. S1. Population dynamics of symbionts in *B. tabaci* MEAM1. Bacterial titers of *Portiera* (*A*) and *Hamiltonella* (*B*) were measured by quantitative PCR in terms of 16S ribosomal RNA gene copies per insect. Each dot represents an individual; filled diamonds, eggs and nymphs; open circles, adult females; gray triangles, adult males; n = 2 for eggs, n = 10 for others. Asterisks indicate statistically significant differences (*P* < 0.001), whereas “ns” indicates not significant (*P* > 0.05). Note that symbiont titer in an egg was calculated by averaging acquired value of 10 individuals.

Fig. S2. *In vivo* localization of facultative symbionts around bacteriocytes in a young adult *B. tabaci*. Female (*A*) and male (*B*) within day 5 after eclosion in MEAM1. Female (*C*) and male (*D*) at day 1 after eclosion in MED Q1. Green indicates *Rickettsia* (*A* and *B*) or *Cardinium* (*C* and *D*). *Portiera* is shown in red. Host nuclear DNA is visualized in blue. Bars, 50 µm.

Fig. S3. *In vivo* localization of *Portiera* (red) and *Hamiltonella* (green) in bacteriocytes of *B. tabaci* MEAM1. (*A*) Adult female at day 1 after eclosion, (*B*) adult male at day 1 after eclosion, (*C*) adult female at day 15 after eclosion, and (*D*) adult male at day 15 after eclosion. N, bacteriocyte nucleus; Bars, 50 µm.

Fig S4. Detection of symbionts and the ER by FISH and immunohistochemistry. The bacteriocyte in a *B tabaci* MEAM1 young adult female at day 1 to 5 after eclosion (*A*), old female at day 15 after eclosion (*B*), old male at day 15 after eclosion (*C*), MED Q1 young adult female at day 1 to 5 after eclosion (*D*), putative young female of *B. tabaci* Asia II 6 (*E*), and *T. vaporariorum* young adult female at day 1 to 5 after eclosion (*F*). The images in *A, D*, and *F* correspond to Fig. 3*A* and *B*, and 5*B*, respectively. Panels for DNA (*first column*), symbiont (*Hamiltonella* or *Arsenophonus*) (*second column*), ER (*third column*), and merged image of the ER and symbiont (*fourth column*) are shown. In (*B, C*, and *F*), panels for *Portiera* are added in the fifth column. DAPI-stained images in the first column show nuclei in the center of bacteriocytes, and *Portiera* and *Hamiltonella* around the nuclei. In merged panels, the ER is shown in violet and *Hamiltonella* or *Arsenophonus* is shown in green. Yellow dashed lines indicate outlines of the bacteriocytes. Bars, 20 µm. Relatively weak signals of *Arsenophonus* and the ER in (*E*) can likely be explained by the use of acetone-preserved samples.

Fig. S5. *In vivo* localization of the ER and endosymbionts in living bacteriocytes of *B. tabaci* MED Q1. Orthogonal view of Z-stack images is shown. Red and blue dashed lines indicate corresponding points in the orthogonal planes. DNA and ER are shown in green and violet, respectively. In the cytoplasm, *Portiera* and *Hamiltonella* were detected as unstructured hypertrophic bacteria (weak green) and rod-shaped bacteria (strong green), respectively. N, nucleus of the bacteriocyte; P, *Portiera*; H, *Hamiltonella*. Bar, 20 µm.

Fig. S6. Interdependent biosynthetic pathways of the essential amino acids in the co-obligate symbionts in whiteflies inferred from the published genomes. Lysine biosynthesis pathways in *(A) B. tabaci* MEAM1 (W. Chen, *et al*. BMC Biol. 14: 110, 2016) and MED Q1 (W. Xie, *et al*. BMC Genomics 19: 68, 2018). Genes of *Portiera, Hamiltonella*, and the host (candidate horizontally transferred genes) are indicated by magenta, green, and blue arrows, respectively. *(B)* Phenylalanine and tryptophan biosynthesis pathways in *T. vaporariorum* (D. Santos-Garcia, C. Vargas-Chavez, A. Moya, A. Latorre, F.J. Silva, Genome Biol. Evol. 7: 873–888, 2015; D. Santos-Garcia, K. Juravel, S. Freilich, E. Zchori-Fein, A. Latorre, A. Moya, S. Morin, F.J. Silva, Front. Microbiol. 9: 2254, 2018). Genes of *Portiera, Arsenophonus* and the host are indicated by magenta, green, and blue arrows, respectively. Missing genes are indicated by dashed arrows. (*C*) Schematic of ER-mediated habitat segregation and putative nutrient flow in the co-obligate symbiotic system in whiteflies.

Fig. S7. Stages of *B. tabaci* MEAM1 development. Yellow cells in the egg or body are bacteriocytes. Numbers of days on the arrow indicate the approximate durations under laboratory conditions. Reproduction is frequently observed in young adults but rarely in senescent adults. Bars, 0.1 mm.

