## Supplemental Movie Legend; Supplemental Figs 1-7; Supplemental Table 1 and 2 for "Subcellular niche segregation of co-obligate symbionts in whiteflies"

Movie S1. Z-stack images of bacteriocytes harboring *Portiera* (red) and *Hamiltonella* (green) in *B. tabaci* MEAM1. Bar, 50 μm. In total, 55 spatially consecutive images were collected by confocal microscopy (depth interval = 0.5 μm) and were processed using Zeiss LSM5 Pascal Image software. Nuclei in the center of bacteriocytes are not shown in the movie to clarify the localization patterns of the two symbionts.

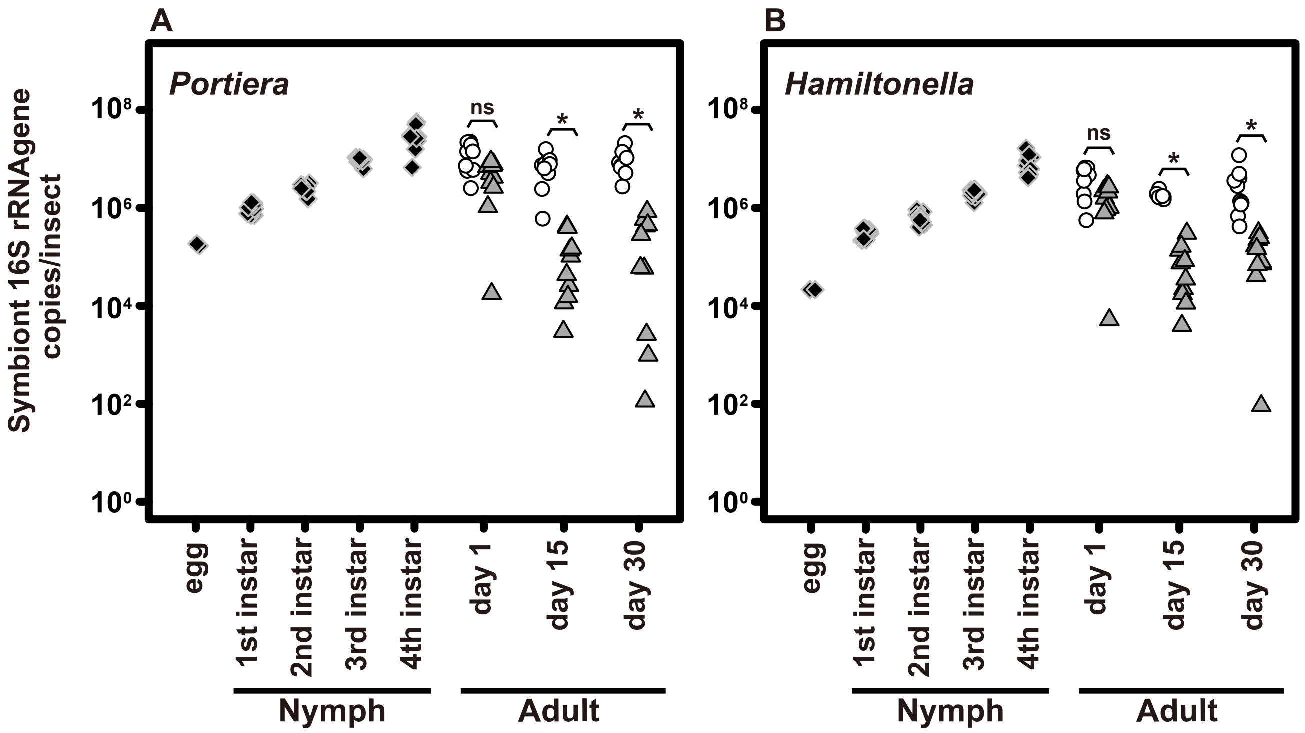


Fig. S1. Population dynamics of symbionts in *B. tabaci* MEAM1. Bacterial titers of *Portiera* (*A*) and *Hamiltonella* (*B*) were measured by quantitative PCR in terms of 16S ribosomal RNA gene copies per insect. Each dot represents an individual; filled diamonds, eggs and nymphs; open circles, adult females; gray triangles, adult males; n = 2 for eggs, n = 10 for others. Asterisks indicate statistically significant differences ($P < 0.001$), whereas “ns” indicates not significant ($P > 0.05$). Note that symbiont titer in an egg was calculated by averaging acquired value of 10 individuals.

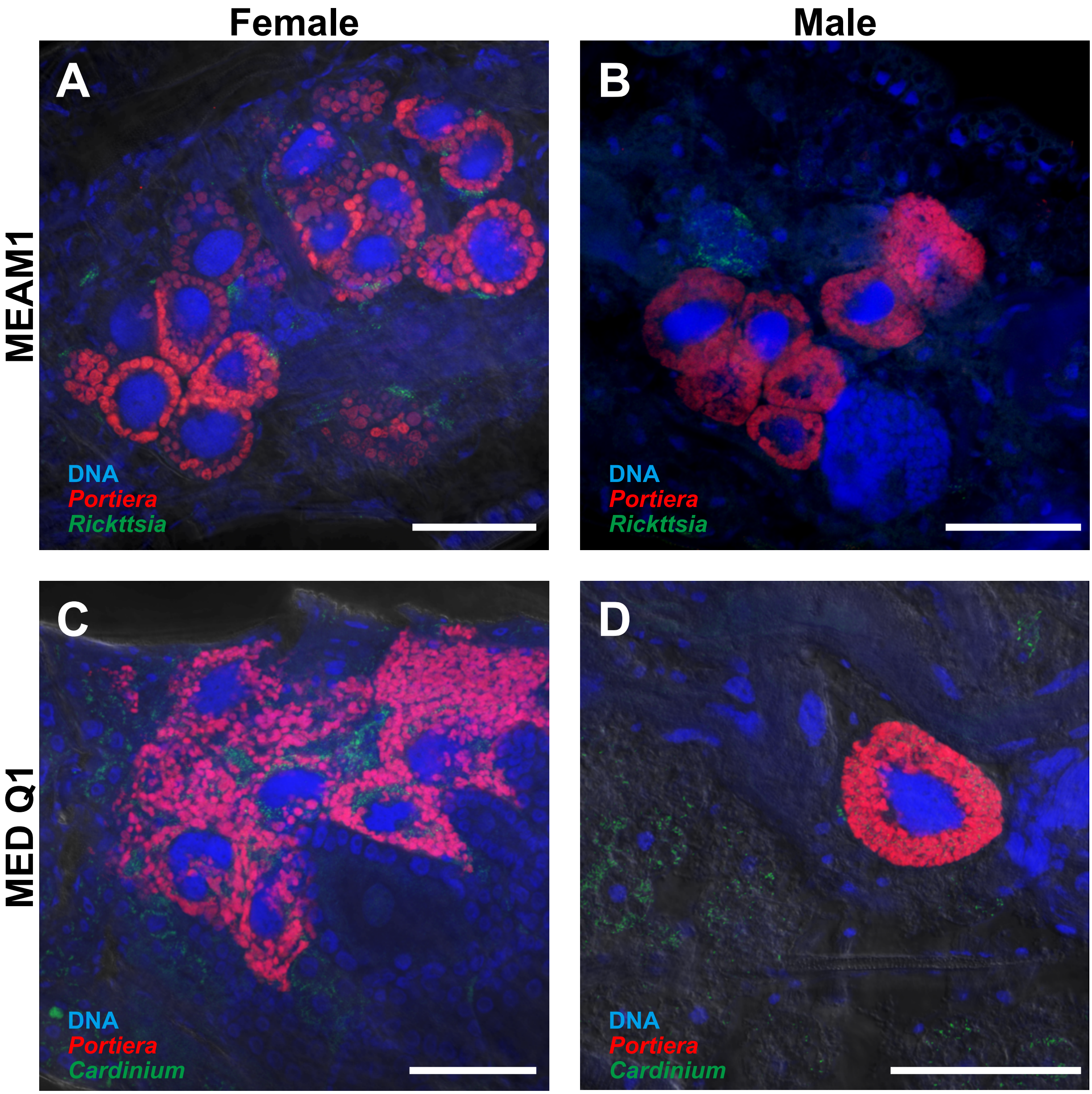


Fig. S2. *In vivo* localization of facultative symbionts around bacteriocytes in a young adult *B. tabaci*. Female (*A*) and male (*B*) within day 5 after eclosion in MEAM1. Female (*C*) and male (*D*) at day 1 after eclosion in MED Q1. Green indicates *Rickettsia* (A and B) or *Cardinium* (*C* and *D*). *Portiera* is shown in red. Host nuclear DNA is visualized in blue. Bars, 50 μm.

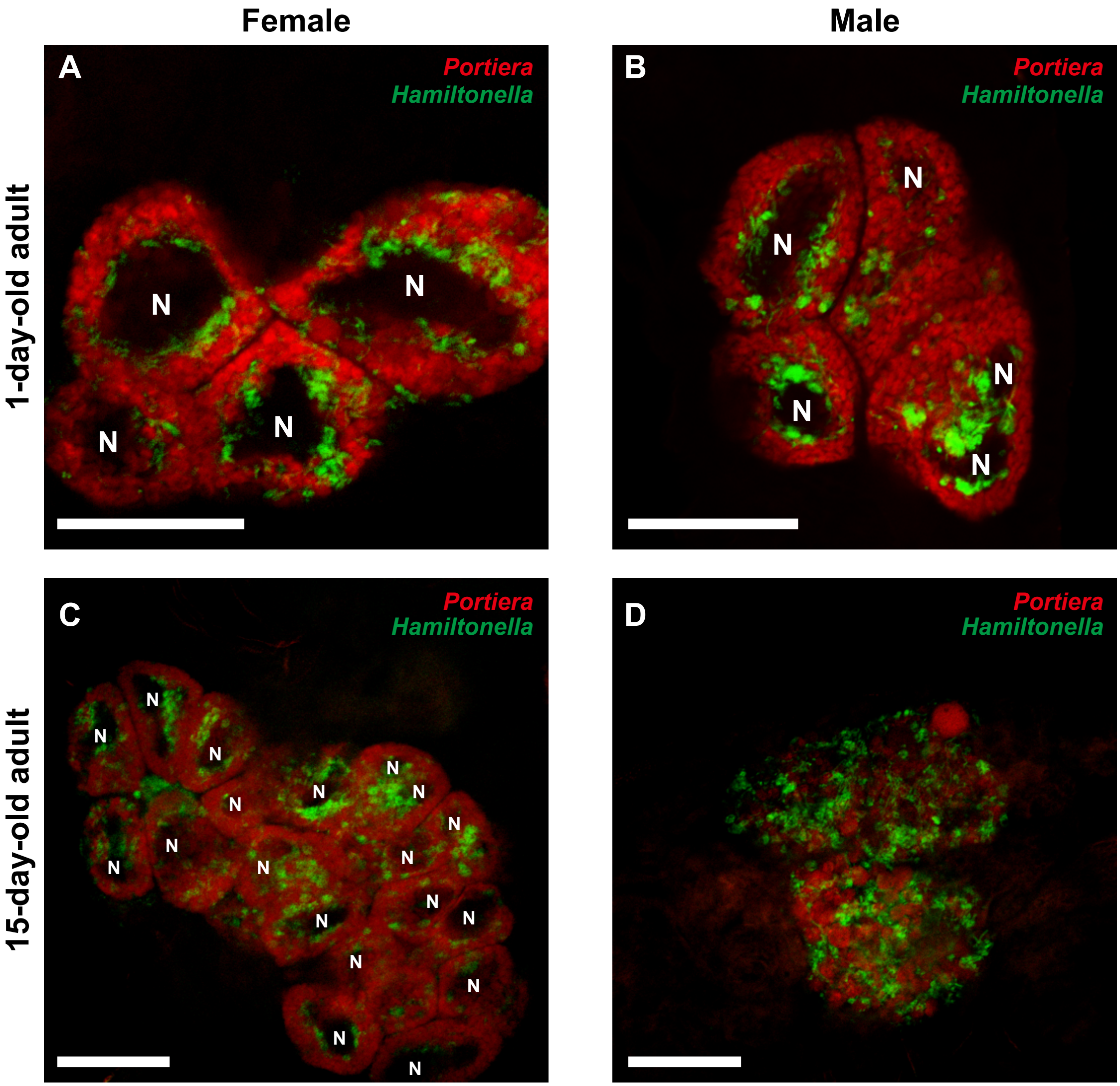


Fig. S3. *In vivo* localization of *Portiera* (red) and *Hamiltonella* (green) in bacteriocytes of *B. tabaci* MEAM1. (*A*) Adult female at day 1 after eclosion, (*B*) adult male at day 1 after eclosion, (*C*) adult female at day 15 after eclosion, and (*D*) adult male at day 15 after eclosion. N, bacteriocyte nucleus; Bars, 50 μm.

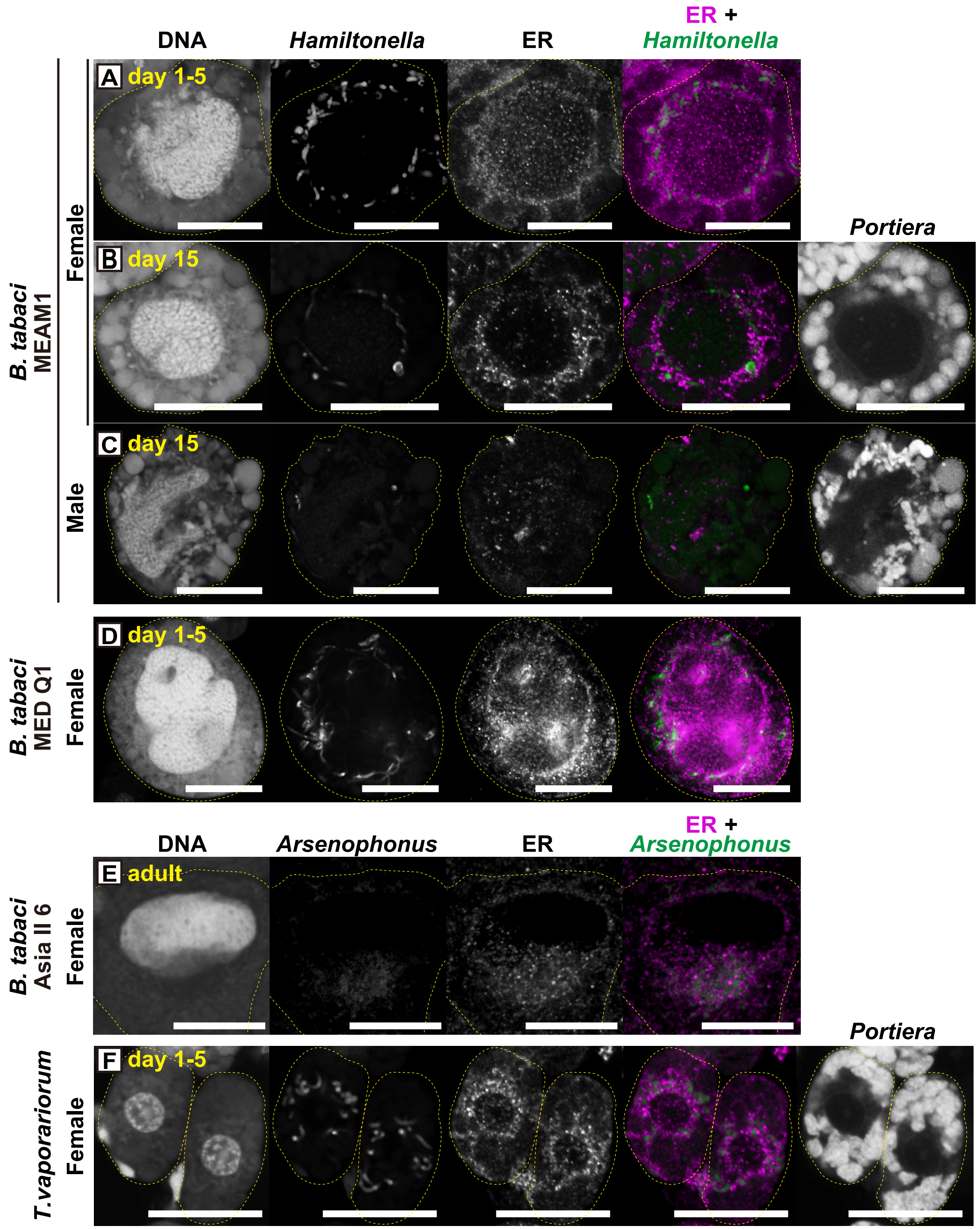


Fig. S4. Detection of symbionts and the ER by FISH and immunohistochemistry. The bacteriocyte in a *B tabaci* MEAM1 young adult female at day 1 to 5 after eclosion (*A*), old female at day 15 after eclosion (*B*), old male at day 15 after eclosion (*C*), MED Q1 young adult female at day 1 to 5 after eclosion (*D*), putative young female of *B. tabaci* Asia II 6 (*E*), and *T. vaporariorum* young adult female at day 1 to 5 after eclosion (*F*). The images in *A*, *D*, and *F* correspond to Fig. 3*A* and *B*, and 5*B*, respectively. Panels for DNA (first column), symbiont (*Hamiltonella* or *Arsenophonus*) (second column), ER (third column), and merged image of the ER and symbiont (fourth column) are shown. In (*B*, *C*, and *F*), panels for *Portiera* are added in the fifth column. DAPI-stained images in the first column show nuclei in the center of bacteriocytes, and *Portiera* and *Hamiltonella* around the nuclei. In merged panels, the ER is shown in violet and *Hamiltonella* or *Arsenophonus* is shown in green. Yellow dashed lines indicate outlines of the bacteriocytes. Bars, 20 µm. Relatively weak signals of *Arsenophonus* and the ER in (*E*) can likely be explained by the use of acetone-preserved samples.

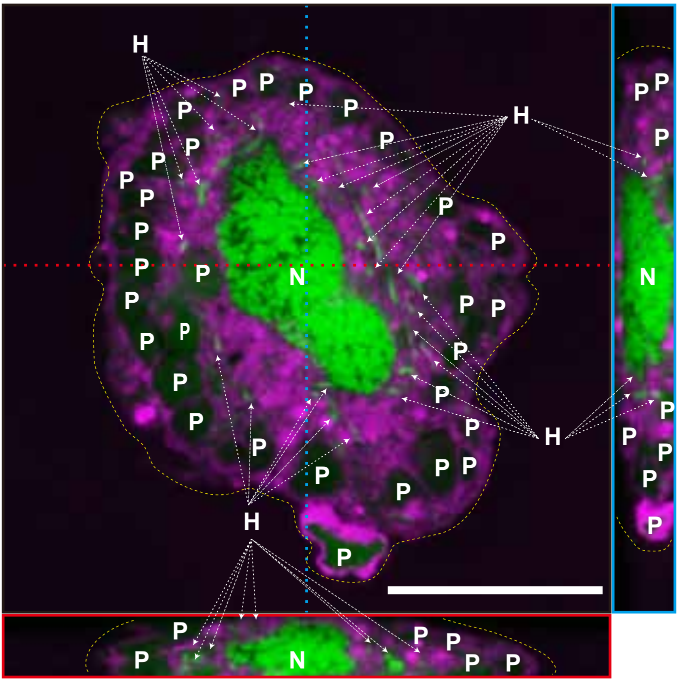


Fig. S5. *In vivo* localization of the ER and endosymbionts in living bacteriocytes of *B. tabaci* MED Q1. Orthogonal view of Z-stack images is shown. Red and blue dashed lines indicate corresponding points in the orthogonal planes. DNA and ER are shown in green and violet, respectively. In the cytoplasm, *Portiera* and *Hamiltonella* were detected as unstructured hypertrophic bacteria (weak green) and rod-shaped bacteria (strong green), respectively. N, nucleus of the bacteriocyte; P, *Portiera*; H, *Hamiltonella*. Bar, 20 μm.

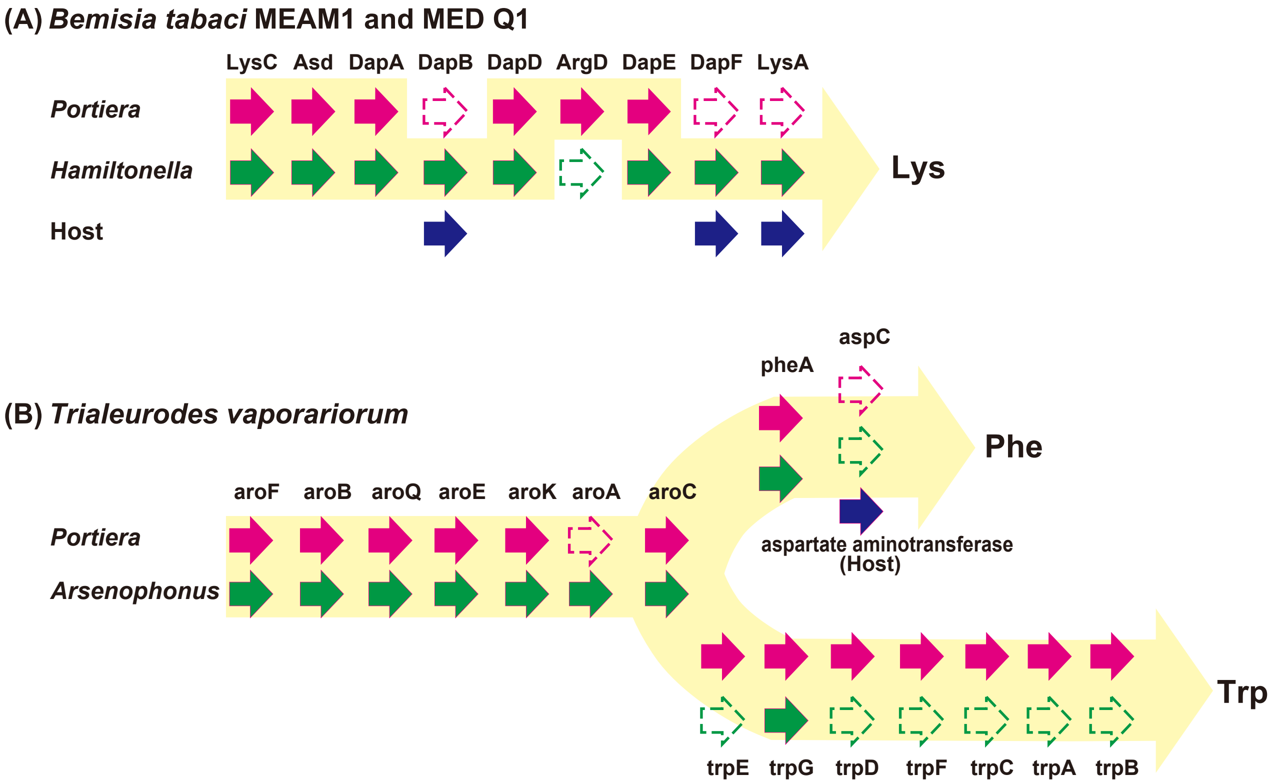


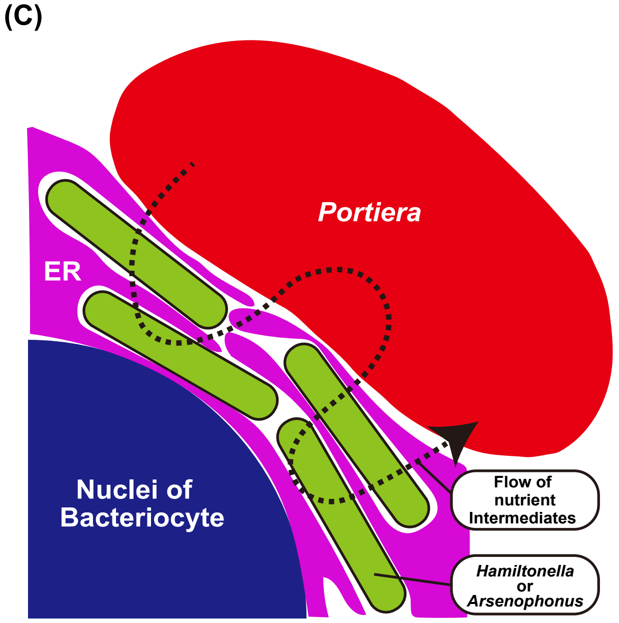


Fig. S6. Interdependent biosynthetic pathways of the essential amino acids in the co-obligate symbionts in whiteflies inferred from the published genomes. Lysine biosynthesis pathways in (A) *B. tabaci* MEAM1 (W. Chen, *et al*. BMC Biol. 14: 110, 2016) and MED Q1 (W. Xie, *et al*. BMC Genomics 19: 68, 2018). Genes of *Portiera*, *Hamiltonella*, and the host (candidate horizontally transferred genes) are indicated by magenta, green, and blue arrows, respectively. (B) Phenylalanine and tryptophan biosynthesis pathways in *T. vaporariorum* (D. Santos-Garcia, C. Vargas-Chavez, A. Moya, A. Latorre, F.J. Silva, Genome Biol. Evol. 7: 873–888, 2015; D. Santos-Garcia, K. Juravel, S. Freilich, E. Zchori-Fein, A. Latorre, A. Moya, S. Morin, F.J. Silva, Front. Microbiol. 9: 2254, 2018). Genes of *Portiera*, *Arsenophonus* and the host are indicated by magenta, green, and blue arrows, respectively. Missing genes are indicated by dashed arrows. (C) Schematic of ER-mediated habitat segregation and putative nutrient flow in the co-obligate symbiotic system in whiteflies.

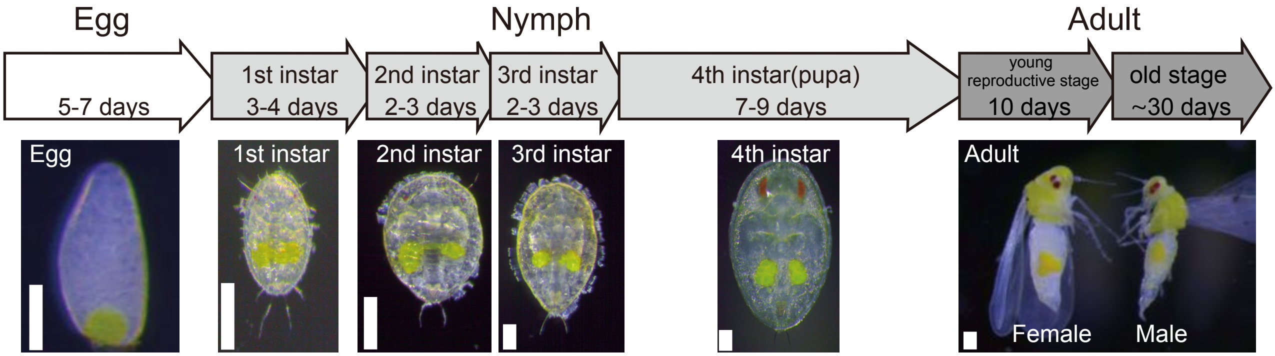


Fig. S7. Stages of *B. tabaci* MEAM1 development. Yellow cells in the egg or body are bacteriocytes. Numbers of days on the arrow indicate the approximate durations under laboratory conditions. Reproduction is frequently observed in young adults but rarely in senescent adults. Bars, 0.1 mm.

**Table S1** Whiteflies used in this study

| Species* | Collection site | Original host plant | Secondary symbionts |
|---|---|---|---|
| *Bemisia tabaci* | | | |
| MEAM1 | Kuwana, Mie prefecture, Japan | *Solanum lycopersicum* | *Hamiltonella, Rickettsia* |
| MED Q1 | Ashikaga, Tochigi prefecture, Japan | *Solanum lycopersicum* | *Hamiltonella, Cardinium* |
| Asia II 6 | Yomitan, Okinawa prefecture, Japan | *Ipomoea batatas* | *Arsenophonus, Wolbachia* |
| *Trialeurodes vaporariorum* | Tsu, Mie prefecture, Japan | *Nicotiana tabacum* | *Arsenophonus* |

* MEAM1 and MED Q1 in *B. tabaci* and *T. vaporariorum* were kept in the laboratory as described in the Materials and Methods section. For Asia II 6, only acetone-preserved samples were used in this study.

**Table S2** PCR primers and oligonucleotide probes used in this study

| Target organism[a] | Primer or Probe sequence (5'→3') | Product size (bp) / fluorechrome | References[d] |
|---|---|---|---|
| **Primers for quantitative PCR** | | | |
| *Portiera* | TGTCAGTGTCAGCCCAGAAG<br>TTTATGTGAAAGCCCTATGCTTA | 157 | 1<br>This study |
| *Hamiltonella* | GCATCGAGTGAGCACAGTTT<br>TATCCTCTCAGACCCGCTAGA | 243 | 2 |
| **FISH probes** | | | |
| *Portiera* in *Bemisia tabaci* | TGTCAGTGTCAGCCCAGAAG | Alexa Fluor 555[b] | 1 |
| *Portiera* in *Trialeurodes vaporariorum* | GCGTACGTTTCCCTACGTTA | Alexa Fluor 555[b] | This study |
| *Hamiltonella* | CCAGATTCCCAGACTTTACTCA | Alexa Fluor 488[b] | 3 |
| *Rickettsia* | TCCACGTCGCCGTCTTGC | Alexa Fluor 488[b] | 1 |
| *Cardinium* | TATCAATTGCAGTTCTAGCG | Alexa Fluor 488[b] | 4 |
| *Arsenophonus* | TCATGACCACAACCTCCAAA | Alexa Fluor 488[b] | 3 |
| **Stellaris RNA FISH probe set** | | | |
| *Hamiltonella* | ctcgacttgcatgtgttagg | Fluorescein[c] | This study |
| | aaactgtgctcactcgatgc | Fluorescein[c] | |
| | ctcgccgacatgaactcagt | Fluorescein[c] | |
| | agattcccagactttactca | Fluorescein[c] | |
| | tttggtctcacgacttcatg | Fluorescein[c] | |
| | ctaccagctaatctcatctg | Fluorescein[c] | |
| | gagatcgtcgcctaggtaag | Fluorescein[c] | |
| | ttcgcccattgtgcaatatt | Fluorescein[c] | |
| | gaaagtgctttacaacccga | Fluorescein[c] | |
| | gtattggcatttatcgcttc | Fluorescein[c] | |
| | tcttctgcgagtaacgtcaa | Fluorescein[c] | |
| | gttattccgattaacgctcg | Fluorescein[c] | |
| | acatctgacttaactcaccg | Fluorescein[c] | |
| | aagttgagctcggggatttc | Fluorescein[c] | |
| | acccagtttcaaatgccatt | Fluorescein[c] | |
| | ccctctagaaaactctagcg | Fluorescein[c] | |
| | caacctccaaatcgacatcg | Fluorescein[c] | |
| | cggacgccacagttcaagac | Fluorescein[c] | |
| | cggtcgatttaacgcgttag | Fluorescein[c] | |
| | tttgagttttaaccttgcgg | Fluorescein[c] | |
| | agttccgtggatgtcaagag | Fluorescein[c] | |
| | tttgtatacgccattgtagc | Fluorescein[c] | |
| | gacgtactttctgagttccg | Fluorescein[c] | |
| | cagactccaatccggactac | Fluorescein[c] | |
| | gacttcatggagtcgagttg | Fluorescein[c] | |
| | aacgtattcaccgtagcatt | Fluorescein[c] | |
| | aagctacctacttcttttgc | Fluorescein[c] | |
| | acaaagtggtaagcgccctc | Fluorescein[c] | |
| | cttcaccccagtcatgaatc | Fluorescein[c] | |
| | aaggaggtgatccaaccgca | Fluorescein[c] | |

**Table S2** ***Continued***

| Target organism[a] | Primer or Probe sequence (5'→3') | Product size (bp) / fluorechrome | References[d] |
|---|---|---|---|
| *Arsenophonus* | ccctatagtgagtcgtatac | Fluorescein[c] | This study |
| | aactctaatcgaattcccgc | Fluorescein[c] | |
| | ctcgacttgcatgtgttacg | Fluorescein[c] | |
| | caaggaagcaagcttccttc | Fluorescein[c] | |
| | caccgtttccagtggttatc | Fluorescein[c] | |
| | cactttgctccttagagatt | Fluorescein[c] | |
| | gaaggtgtgaggccagaacg | Fluorescein[c] | |
| | gctaatctcatatgggttca | Fluorescein[c] | |
| | tagagatcgtcgcctaggtg | Fluorescein[c] | |
| | caccttcctcacgactgaaa | Fluorescein[c] | |
| | ttgctaaggttattaaccct | Fluorescein[c] | |
| | tcttctgtcgctaacgtcaa | Fluorescein[c] | |
| | ttaattaaccgcctgcgtgc | Fluorescein[c] | |
| | ccggggatttcacattcaac | Fluorescein[c] | |
| | accagtcttgaatgccattc | Fluorescein[c] | |
| | cctctacaagactctagcta | Fluorescein[c] | |
| | tacatatggaattctacccc | Fluorescein[c] | |
| | cacatgagcgtcagtctttg | Fluorescein[c] | |
| | atgaccacaacctccaaatc | Fluorescein[c] | |
| | gtcgatttacgcgttagctc | Fluorescein[c] | |
| | tttgagttttaaccttgcgg | Fluorescein[c] | |
| | gattcgctggatgtcaagag | Fluorescein[c] | |
| | gcactcctctatctctaaag | Fluorescein[c] | |
| | gataagggttgcgctcgttg | Fluorescein[c] | |
| | ccgaatcgctggcaacaaag | Fluorescein[c] | |
| | ccatgatgacttgacgtcat | Fluorescein[c] | |
| | ctctgtatacgccattgtag | Fluorescein[c] | |
| | gacgtactttctgagttccg | Fluorescein[c] | |
| | cagactccaatccggacttc | Fluorescein[c] | |
| | gacttcatggagtcgagttg | Fluorescein[c] | |
| | aacgtattcaccgcgacatg | Fluorescein[c] | |
| | aagctacctacttcttttgc | Fluorescein[c] | |
| | tggtaagcgccatccaaaaa | Fluorescein[c] | |

[a] Taget gene is 16S ribosomal RNA. [b] The probe is labelled at the 5' end. [c] The probe is labelled at the 3' end. [d] 1, Y. Gottlieb, M. Ghanim, E. Chiel, D. Gerling, V. Portnoy, S. Steinberg, G. Tzuri, A.R. Horowitz, E. Belausov, N. Mozes-Daube, S. Kontsedalov, M. Gershon, S. Gal, N. Katzir, E. Zchori-Fein, Identification and localization of a *Rickettsia* sp. in *Bemisia tabaci* (Homoptera: Aleyrodidae). Appl. Environ. Microbiol. 72: 3646–3652 (2006); 2, M. Brumin, S. Kontsedalov, M. Ghanim, *Rickettsia* influences thermotolerance in the whitefly *Bemisia tabaci* B biotype. Insect Sci. 18: 57–66 (2011); 3, Y. Gottlieb, M. Ghanim, G. Gueguen, S. Kontsedalov, F. Vavre, F. Fleury, E. Zchori-Fein, Inherited intracellular ecosystem: symbiotic bacteria share bacteriocytes in whiteflies. FASEB J. 22: 2591–2599 (2008); 4, M. Skaljac, K. Zanic, S.G. Ban, S. Kontsedalov, M. Ghanim, Co-infection and localization of secondary symbionts in two whitefly species. BMC Microbiol. 10: 142 (2010).
